# Nuclear Myosin 1 regulates platelet activation and immune response: A novel therapeutic target for hematologic disorders

**DOI:** 10.1101/2023.02.14.528461

**Authors:** Tomas Venit, Samira Khalaji, Valentina Fambri, Wael Abdrobou, Mei El Gindi, Rahul Shrestha, Pavel Hozak, Youssef Idaghdour, Jeremy CM Teo, Piergiorgio Percipalle

## Abstract

Cellular differentiation involves complex events associated with changes in cellular shape, function and proliferative capacity. This process is regulated by specific expression of multiple genes, which guide the cell through differentiation but also ensure proper function of terminal cell types. Over the last decade, the role of cellular metabolism in maintaining stem cells pluripotency and differentiation has been getting more attention due to a direct link between metabolic state and differentiation potential of cells. Since Nuclear Myosin 1 (NM1) deletion leads to a switch from oxidative phosphorylation to aerobic glycolysis and tumorigenesis in mice, here we asked if NM1 also contributes to cell differentiation. Indeed, analysis of metabolomic profiles and cytokine and chemokine levels upon NM1 depletion support a role in hematopoiesis. In an NM1 KO mouse model results from total blood cell count and bleeding assay show decreased erythropoiesis and thrombopoiesis as well as failure in hemostasis. Next, we explored the role of NM1 during the differentiation of hematopoietic progenitor stem cells to terminal blood cells by transcriptionally profiling bone marrow, spleen and peripheral blood from NM1 KO mice. We found that in bone marrow, NM1 deletion leads to overexpression of genes associated with glycolysis-dependent platelet activation and suppression of the innate immune system. These expression patterns are preserved and become more complex in spleen and peripheral blood. The study, therefore, provides insights into the underlying mechanisms of hematopoietic differentiation and activation of specific blood cell types and suggests NM1 as a potential therapeutic target for blood-related disorders.

## Introduction

Nuclear Myosin 1 (NM1) belongs to the family of unconventional single-headed myosins present in the cytoplasm and the cell nucleus. Myosins are involved in various processes such as regulation of the plasma membrane and actin cytoskeleton, exocytosis of vesicles, cell movement, repair of DNA damage foci, movement of the chromatin, and transcription regulation. NM1 was the first myosin discovered in the cell nucleus, and over the years, several studies described its general roles and mechanism of function in Polymerase I, II, and III transcription (1, 2). NM1 is one of three alternatively spliced isoforms of the Myo1C gene exhibiting an extra 16 amino-acid stretch on its N-terminus. All Myo1C isoforms share the same nuclear localization signal and partially overlap in function (3, 4). In the cell nucleus, NM1 has been shown to directly bind to the chromatin at the transcription start site, forming a complex with actin and the polymerase machinery, which is later on released and replaced by NM1 binding to the chromatin remodeling complex B-WHICH via direct interaction with the ATPase subunit SNF2H (5, 6). This leads to chromatin remodeling around the transcription start site but also to NM1 association with the histone-acetyl- and histone-methyl-transferases PCAF and Set1, respectively. They acetylate and methylate surrounding histones as a mark for active transcription and allow polymerase-mediated transcription. Recently, we reported that these mechanisms are important for numerous cellular functions, including the transcriptional response to DNA damage (7). Mechanistically, we discovered that NM1 is part of the DNA damage signaling cascade as it interacts with p53 to activate transcription of the cell cycle regulator p21 (Cdkn1A). Upon DNA damage, p53 binds to its responsive promoter upstream of the Cdkn1A gene followed by NM1-dependent acetylation and methylation of surrounding H3 histones. This allows for rapid p21 protein accumulation and cell cycle arrest until DNA damage is repaired, while cells missing Nuclear myosin 1 accumulate DNA damage and are prone to apoptosis (2, 7).

In a more recent study, we reported a novel role for NM1 as a tumour suppressor and regulator of cellular metabolism. The conventional view of somatic cells using only oxidative phosphorylation and cancer cells using only aerobic glycolysis as a primary type of metabolism was widely disproved over time. In many cases, both types of metabolism coexist in cells, with their ratio adjusting based on the acute needs. For example, pluripotent stem cells, in many ways resembling cancer cells, use aerobic glycolysis as the primary energy source, but upon differentiation, they gradually move towards oxidative phosphorylation, which becomes a dominant metabolic pathway. However, some fully differentiated somatic cells can still rely on aerobic glycolysis as their primary energy source or switch between the two metabolic pathways depending on the surrounding environment. We showed that NM1 is part of the nutrient-sensing PI3K/Akt/mTOR pathway, forming a positive feedback loop with mTOR. NM1 deletion leads to a suppression of mitochondrial transcription factors TFAM and PGC1α, negatively regulating mitochondrial oxidative phosphorylation and leading to metabolic reprogramming towards aerobic glycolysis associated with cancer cells. Indeed, NM1 knockout (KO) cells can form solid tumours in a nude mouse model even though they have suppressed the PI3K/Akt/mTOR signalling pathway, suggesting that the metabolic switch towards aerobic glycolysis provides a sufficient signal for carcinogenesis (8).

Seeing that NM1 regulates cellular metabolism by influencing the transcription of mitochondrial transcription factors, in this study, we investigated whether NM1 deletion affects the ability of pluripotent stem cells to differentiate into terminal somatic cells. Additionally, we examined whether NM1 impacts the functionality of specific cell types that rely on active switching between two metabolic pathways. To address this, we used hematopoiesis as a model system since hematopoietic stem cells (HSCs) give rise to a wide variety of highly specialized blood cells with specific energetic needs. In general, these cells are present in the bone marrow, and they are generated from hematopoietic lineages with progenitor hematopoietic stem cells (HSCs) giving rise to red blood cells, white blood cells, and platelets. All other cell types are components of the bone marrow stroma and are involved in the formation of specific microenvironments regulating HSCs. The cell types present in bone marrow stroma include mesenchymal stem cells, fibroblasts, adipocytes, osteoblasts, osteoclasts, and endothelial cells (9). From the metabolic perspective, hematopoietic stem cells use anaerobic glycolysis as the only energy source and switch to oxidative phosphorylation only during differentiation (10). The energy metabolism of fully differentiated hematopoietic cells depends on their function. Erythrocytes lose their nucleus and mitochondria during differentiation, making them entirely reliant on glycolysis for energy production (11). Monocytes use oxidative phosphorylation as the primary energy source. However, they switch to glycolysis upon activation to pro-inflammatory M1 macrophages and back to oxidative phosphorylation upon switching to anti-inflammatory M2 macrophages (12, 13). Like monocytes, lymphocytes use oxidative phosphorylation in their quiescent state and switch to glycolytic metabolism only upon activation (14). In contrast, neutrophils use glycolysis, and their limited mitochondria serve only for apoptotic signaling (15). Finally, platelets (thrombocytes), derived from the cytoplasmic fragmentation of megakaryocytes and essential for blood coagulation at injury sites, utilize both oxidative phosphorylation and glycolysis for energy production. However, their activation depends on aerobic glycolysis (16, 17). The bone marrow stromal cells follow similar metabolic patterns as hematopoietic lineages, and mesenchymal stem cells use glycolysis as a primary source of metabolism (18). Differentiation of these cells to osteoblasts, however, is not associated with metabolic switch to oxidative phosphorylation, and osteoblasts use glycolysis to produce ATP (19). In contrast, osteoclasts, which originate from hematopoietic monocytes, rely on oxidative phosphorylation during differentiation and maturation but switch to glycolytic metabolism only upon activation (20, 21).

By using NM1 wild type (WT), knock-out (KO) and NM1-rescued (KO+NM1) mouse embryonic fibroblasts (MEFs), as well as NM1 WT and KO mice, here we show evidence that NM1 deletion leads to alterations in signature metabolites implicated in hematopoiesis and differential gene expression of genes involved in platelet activation and immune system response. In an NM1 KO mouse we show decreased erythropoiesis and thrombopoiesis associated with failure in hemostasis. Consistent with our previous results, showing the shift towards glycolysis in NM1 KO cells, we show that glycolysis-dependent platelet activation is upregulated upon NM1 deletion in bone marrow and peripheral blood. In contrast, the innate immune system response is suppressed in NM1 KO mice, which can be associated with the need for oxidative phosphorylation during differentiation of several innate immune cells, like monocytes, dendritic cells, or mast cells. In agreement with it, bead-based immunoassays show that the excretion of several cytokines and pro-inflammatory chemokines is abolished in NM1 KO cells. In the spleen, innate immune system genes are suppressed as well, while adaptive immune system genes, mostly associated with T-cell activation and T-regulatory cell (Treg) differentiation, are upregulated, which is consistent with the metabolic need for aerobic glycolysis in these cells.

### Material and Methods Cell lines

Nuclear Myosin 1 Knock-Out (NM1 KO) cell lines were derived from wild-type mouse embryonic fibroblasts (MEFs) (ATCC® CRL-2752) (NM1 WT) using the CRISPR/Cas9 system (7). The rescue cell line expressing exogenous NM1 in NM1 KO MEFs (NM1 KO+NM1) was prepared previously by lentiviral transduction of vector carrying coding sequence for NM1 fused with HA and V5 tag (VectorBuilder) (8). Actin WT and KO cells were described previously (22, 23). Cells were grown in a DMEM medium containing 10% fetal bovine serum, 100 U/ml penicillin, and 100 mg/ml streptomycin (Millipore-Sigma) in a humidified incubator with 5% CO_2_ at 37 °C.

### Large-scale Ultra-Performance Liquid Chromatography High-Resolution Mass Spectrometry

5 replicates of 3x10^5^ overnight grown NM1 WT and KO cells were used for metabolomic profiling. Cells were washed twice with ice-cold 0.9% NaCl solution and then incubated on ice for 3 minutes in 300µl of 100% Methanol. Subsequently, cells were scraped using a precooled cell scraper, moved to a cold Eppendorf tube, and spun down at 15000rpm for 15 min at 4°C. 200µl of each supernatant was used for further processing. Samples were vacuum dried and reconstituted in 200µl of cyclohexane/water (1:1) solution for subsequent analysis by Ultra Performance Liquid Chromatography High-Resolution Mass Spectrometry (UPLC-HRMS) performed at the VIB Metabolomics core facility (Belgium). 10 ul of each sample were injected on a Waters Acquity UHPLC device connected to a Vion HDMS Q-TOF mass spectrometer. Chromatographic separation was carried out on an ACQUITY UPLC BEH C18 (50 × 2.1 mm, 1.7 μm) column from Watersunder under the constant temperature of 40°C. A two-buffer gradient was used for separation: buffer A (99:1:0.1 water:acetonitrile:formic acid, pH 3) and buffer B (99:1:0.1 acetonitrile:water: formic acid, pH 3), as follows: 99% A for 0.1 min decreased to 50% A in 5 min, decreased to 30% from 5 to 7 minutes, and decreased to 0% from 7 to 10 minutes. The flow rate was set to 0. 5 mL min−1. Both positive and negative Electrospray Ionization (ESI) were applied to screen for a broad array of chemical classes of metabolites present in the samples. The LockSpray ion source was operated in positive/negative electrospray ionization mode under the following specific conditions: capillary voltage, 2.5 kV; reference capillary voltage, 2.5 kV; source temperature, 120°C; desolvation gas temperature, 600°C; desolvation gas flow, 1000 L h−1; and cone gas flow, 50 L h−1. The collision energy for the full MS scan was set at 6 eV for low energy settings, for high energy settings (HDMSe) it was ramped from 28 to 70 eV. The mass range was set from 50 to 1000Da, scan time was set at 0.1s. Nitrogen (greater than 99.5%) was employed as desolvation and cone gas. Leucine-enkephalin (250 pg/μL solubilized in water: acetonitrile 1:1 [v/v], with 0.1% formic acid) was used for the lock mass calibration, with scanning every 1 min at a scan time of 0.1 s. Profile data was recorded through Unifi Workstation v2.0 (Waters).

### Unsupervised Metabolomics Data Analysis

Raw MS peak intensity data representing the abundance of detected metabolic features were subject to data integrity check. Metabolic features with a constant or single value across all samples were removed. Missing values, if detected, were replaced by 1/5 of the minimum positive values of their corresponding variables. Data filtering was performed to remove non-informative features that are considered baseline noise. Features that are near-constant throughout experimental conditions were detected using standard deviation (SD) and removed from downstream analysis (24). Filtered data were subject to sample median normalization to adjust for systematic differences among samples. Data were then log-transformed (base 10) to deal with negative values after the normalization. Pareto scaling was used to adjust each metabolic feature by a scaling factor computed based on the dispersion of the variable (mean-centered and divided by the range of each variable). PCA and hierarchical clustering were done to explore the correlation structure in the data across the two experimental conditions (WT and KO). Data QC, filtering, and unsupervised statistical analysis were performed using MetaboAnalyst v6.0 (25). An unpaired t-test was used to detect differential abundant metabolic features between the experimental conditions.

### Functional and metabolic pathway enrichment analysis

Functional analysis of curated normalized peak intensity data was performed using MetaboAnalyst v6.0 using an existing protocol (25). Implemented Gene Set Enrichment Analysis (GSEA) method in the Functional Analysis module of MetaboAnalyst v6.0 (Accessed in 2024 from http://www.metaboanalyst.ca/) was used to identify sets of functionally related compounds and evaluate their enrichment of potential functions defined by metabolic pathways. GSEA analysis of compounds identified using positive and negative ionization was performed separately. Putative annotation of MS peaks data considering different adducts and ion modes was performed. *m*/*z* values and retention time dimensions were used to increase confidence in identifying compounds and improve the accuracy of functional interpretations. Annotated compounds were then mapped onto Mus musculus (mouse) [BioCyc] (26–28) for pathway activity prediction (Supplementary Table 1 and 2). GSEA calculates the Enrichment score (ES) by walking down a ranked list of metabolites, increasing a running-sum statistic when a metabolite is in the metabolite set and decreasing it when it is not. A metabolite set is defined in this context as a group of metabolites with collective biological functions or common behaviors, regulations, or structures. In this method, the ES of each enriched pathway is calculated to reflect the degree to which a metabolite set is overrepresented at the top or bottom of a ranked list of metabolites. Each ES is then normalized by the average of all ES scores against all permutations of the expression dataset to generate normalized enrichment scores (NES) that are used to compare analysis results across metabolite sets. By normalizing the enrichment score, GSEA accounts for differences in metabolite set size and correlations between metabolite sets and the expression dataset. A positive NES indicates that the enriched metabolite set is at the top of the ranked list; a negative NES indicates that the enriched metabolite set is at the bottom of the ranked list (28). Finally, compound hits were identified for each enriched pathway. Raw metabolomic data are publicly available in the Mendeley database (https://data.mendeley.com/preview/nxzs4dtztg?a=aecce665-2b58-4a98-8328-b70d689c249a).

### Chemokine and cytokine measurement

NM1 WT, NM1 KO, and NM1 KO+NM1 were used for the analysis. 5x 10^5^ cells per well were seeded in 24 well plates and left to grow for 48 hours under standard conditions. Next, cells were spun down on plates to remove debris and media used for subsequent analysis of chemokine and cytokine content by the bead-based immunoassays LEGENDplexTM Mouse Cytokine Panel 2 (740134) and LEGENDplexMU Proinflammatory Chemokine Panel (both BioLegend) according to manufacturer’s protocol. In short, 25µl of media from each cell type was incubated for 2 hours with a combination of different capture beads bound to specific antibodies detecting given chemokine/cytokine. After a series of washes, biotinylated detection antibodies were added to each sample plate to form capture bead-analyte-detection antibody sandwiches. Next, streptavidin-phycoerythrin (SA-PE) was added to the mixture to be bound to the biotinylated detection antibodies, providing fluorescent signal intensities in proportion to the number of bound analytes. Samples were then analyzed by flow cytometer, where specific beads were differentiated by size and internal fluorescence intensity, and analyte-specific populations were segregated and quantified by the PE fluorescent signal. The concentration of a particular analyte was determined by a standard curve generated in the same assay. For each cell type, 6 biological replicates were used for the measurement and each measurement was performed in 2 technical replicates. The final concentration of the given analyte in each biological replicate is calculated as a mean value between 2 technical replicates.

### Isolation of tissues from mice and isolation of RNA

Wild type (WT) and NM1 KO C57/Bl6 mice (8 months old; 25-35 g in weight) were used for the analysis (29). The mice were housed in a pathogen-free sterile ventilation system supplied with sterile woodchip and *ad libitum* feed and water supply. Mice were kept in a room maintained at a temperature of 21-24 °C on a schedule of 12 h light/dark cycle. All animal experiments were performed after approval by the NYUAD-IACUC (Protocol 20-0004, 23-0009).

For the transcriptomic analysis we used bone marrow tissue as an example of the primary lymphoid organ where hematopoietic differentiation happens, the spleen as an example of a secondary lymphoid organ where specific mature cell types reside, and are activated upon stimulus, and peripheral blood as the site of action for circulating blood cells. Bone marrow was isolated according to the previously published protocol (30). In short, femurs and tibiae were taken from euthanized mice and cleaned from surrounding muscles, connective tissues, and epiphysis parts of each long bone. All bones from single mice were placed in the 0.5ml tube with the hole on the bottom nested in the 1.5ml collection tube. Bones were spun down at 14000 x g in a microcentrifuge for 15 seconds to pellet the bone marrow at the bottom of the bigger tube. For each bone marrow sample, femurs and tibiae from single NM1 WT or KO mice were used. RNA was isolated by RNAzol according to the manufacturer’s protocol. Spleen was isolated from euthanized mice and cleaned from surrounding fat and connective tissue. Approximately 5mm^3^ of tissue from the tip of the spleen was merged in RNAzol and homogenized in bead mill homogenizer followed by standard RNA isolation protocol. Blood samples were collected by submandibular vein bleeding according to a previously published protocol (31). Collected blood was immediately mixed with the double amount of RNAzol and samples were thoroughly mixed and stored on dry ice before RNA isolation by standard protocol. The final RNA concentration was measured by the Qubit RNA BR Assay Kit.

### RNA-Seq library preparation, sequencing, and analysis

Three replicates were prepared for each tissue and each genotype. 500ng of total RNA was used for RNA-Seq library preparation with the NEBNext Ultra II RNA Library Prep Illumina Kit (NEB) and sequenced with the NextSeq 500/550 sequencing platform (performed at the NYUAD Sequencing Center within the NYUAD Core Technology Platform). Subsequent analysis, including quality trimming, was executed using the BioSAILs workflow execution system. Trimmomatic (version 0.36) was used for quality trimming of the raw reads to get rid of low-quality bases, systematic base-calling errors, and sequencing adapter contamination(32). The quality of the sequenced reads pre/post quality trimming was assessed by FastQC and only the reads that passed quality trimming in pairs were selected for downstream analysis (https://www.bioinformatics.babraham.ac.uk/projects/fastqc/). The quality-trimmed RNA-Seq reads were aligned to the Mus musculus GRCm38 (mm10) genome using HISAT2 (version 2.0.4)(33). The conversion and sorting of SAM alignment files for each sequenced sample to BAM format were done by using SAMtools (version 0.1.19)(34). The BAM alignment files were processed using HTseq-count, using the reference annotation file to produce raw counts for each sample. The raw counts were analyzed using the NASQAR online analysis tool (http://nasqar.abudhabi.nyu.edu/), to merge, normalize, and identify differentially expressed genes (DEG) as well as for the production of PCA and MA plots. DEG with log2(FC) ≥ 0.5 and adjusted p-value of <0.05 for upregulated genes, and log2(FC) ≤ −0.5 and adjusted p-value of <0.05 for downregulated genes between the NM1 WT and KO samples for each tissue were used for the gene ontology (GO) enrichment analysis using DAVID Bioinformatics (https://david.ncifcrf.gov/)(35). RNA-Seq data were deposited in the Gene Expression Omnibus (GEO) repository under accession number GSE293993. Gene ontology bar charts and gene expression heatmaps were produced in GraphPad Prism 8.3.0 software. The STRING database was used for the production of an interaction map based on text mining, experimental data, co-expression, and database search (36).

### Bleeding Time (BT) Test

To assess primary hemostasis and to investigate the effect of NM1 on platelet function, bleeding time was measured following tail-tip transection. Briefly, 2 mm of the distal tail tip was amputated, and the bleeding site was gently blotted every 15 seconds using the rough side of filter paper without applying pressure to the wound. The procedure continued until no further blood appeared on the filter paper. Bleeding time (in seconds) was calculated by multiplying the total number of blood spots by 15. The physiological bleeding time range in mice typically falls between 2 to 7 minutes (37).

### Complete Blood Count (CBC)

Whole blood was collected from 12-month-old mice via cardiac puncture into EDTA-coated tubes to prevent coagulation. Haematological parameters were analysed using an automated BeckmanDxH 900 Haematology Analyzer according to the manufacturer’s instructions. Platelet count (PTL), white blood cell count (WBC), red blood cell count (RBC), and unclassified white blood cells (UWBC) were measured. Differential WBC counts, including neutrophils, lymphocytes, monocytes, eosinophils, basophils, and nucleated red blood cells (NRBC), were also recorded. Additional parameters, including hematocrit (HCT), hemoglobin concentration (HGB), mean corpuscular volume (MCV), mean corpuscular hemoglobin (MCH), mean corpuscular hemoglobin concentration (MCHC), red cell distribution width (RDW and RDW-SD), and mean platelet volume (MPV), were evaluated to assess platelet numbers and overall haematological profiles in both WT and KO mice.

## Results

### Hematopoietic signature metabolites are dysregulated in NM1-depleted mouse embryonic fibroblasts

To study the impact of NM1 on metabolic processes, we performed Liquid Chromatography followed by High-Resolution Mass Spectrometry of cellular extracts from WT and NM1 KO MEFs. A total of 15,423 features were detected, and the quality control of the peak intensity data resulted in the retention of 9,193 and 5,425 metabolic features detected using positive and negative ionization, respectively.

Principal-component analysis (PCA) of the normalized peak intensity data detected using positive ionization revealed a strong correlation structure in the metabolomic data across both conditions with the first two PCs explaining 66.0% of the variation in the dataset (Supplementary Figure 1A). Similarly, PCA of normalized peak intensity data detected using negative ionization revealed a strong correlation structure in the metabolomic data across both conditions with the first two PCs explaining 61.5% of the variation in the negative ionization dataset (Supplementary Figure 1B), capturing the effect of loss of NM1 and showing clear segregation between replicates of the WT and KO. Next, we used an unpaired t-test to investigate differentially abundant metabolic features between NM1 WT and KO cells. Clustering of the top 50 significantly differentially abundant metabolic features between the two conditions clearly shows that replicates of each condition cluster together and NM1 deletion leads to distinct metabolomic changes in these cells in both datasets using the two ionization methods (Supplementary Figure 1C and 1D).

To identify perturbations in mouse-specific metabolic pathways or metabolite sets in association with the loss of NM1, functional analysis using the Gene Set Enrichment Analysis (GSEA) approach was performed for metabolic features detected using positive and negative ionization and mapped onto Mus musculus (mouse) [BioCyc], separately. The analysis of the positive ionization dataset revealed the enrichment of polyamine biosynthesis pathways (Spermine biosynthesis I and II, and Spermidine biosynthesis pathways, NES score = 1.281, *p-value* = 0.04; Supplementary Table 3 and Figure 1A). Particularly, the analysis revealed a significant increase in the abundance of 5-methylthioadenosine (MTA) in the NM1 KO Cells (*t* = 6.55, *p* value = 0.0002; Supplementary Table 1 and 3, and Figure 1C). The significant increase in the levels of MTA, the S-methyl derivative of adenosine, in the NM1-KO cells reveals significant perturbations in the methionine salvage pathway, which is universal to aerobic life (38, 39). Importantly, MTA, a naturally occurring nucleoside, plays a significant role in modulating platelet aggregation. It has been shown that MTA and its analogs regulate platelet aggregation through its influence on adenosine signaling pathways. MTA has been shown to elevate intracellular cAMP levels, which subsequently inhibits platelet aggregation. This mechanism operates independently of the P2Y12 receptor, a key player in platelet activation, suggesting that MTA interacts with an unidentified G protein-coupled receptor that stimulates cAMP production (40, 41).

**Figure 1.**
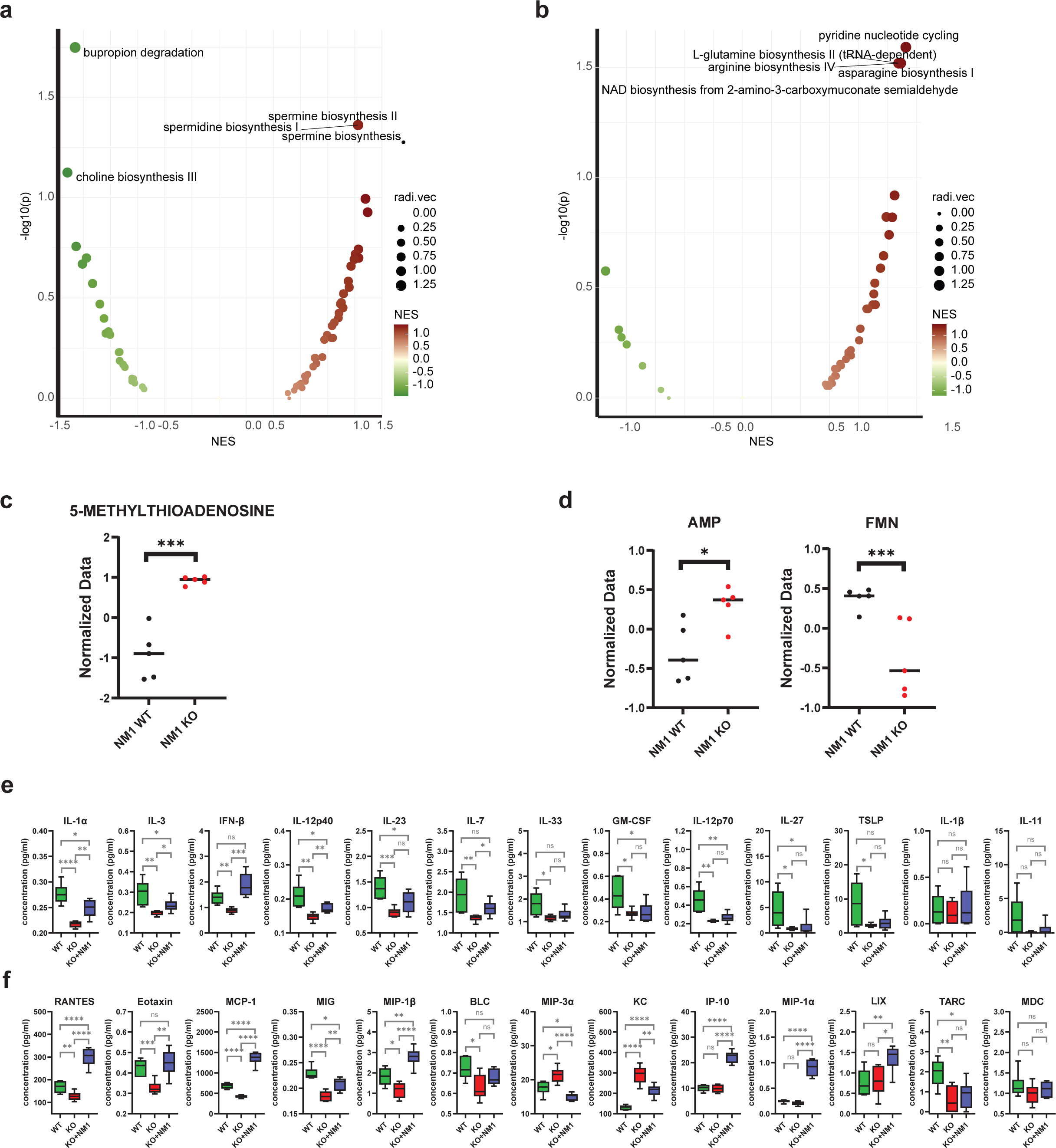
NM1 deletion leads to changes in hematopoietic signature metabolites and cytokine/chemokine production in mouse embryonic fibroblasts. **(a)** Gene Set Enrichment Analysis (GSEA) for metabolic features detected using positive ionization. **(b)** Gene Set Enrichment Analysis (GSEA) for metabolic features detected using negative ionization. **(c)** Metabolomic profile of 5-methylthioadenosine (MTA) across experimental samples as detected by positive ionization. *** P≤0.001 **(d)** Metabolomic profile of AMP and FMN across experimental samples as detected by negative ionization. * P≤0.05, *** P≤0.001 **(e)** List of analyzed cytokines in NM1 WT, NM1 KO and rescued NM1 KO+NM1 cells. ns P > 0.05, * P≤0.05, ** P≤0.001, *** P≤0.001, **** P≤0.0001. **(f)** List of analyzed chemokines in NM1 WT, NM1 KO, and rescued NM1 KO+NM1 cells. ns P > 0.05, * P≤0.05, ** P≤0.001, *** P≤0.001, **** P≤0.0001.

The analysis of the negative ionization dataset revealed the enrichment of pyridine nucleotide cycling, L-glutamine biosynthesis II (tRNA-dependent), asparagine biosynthesis I, arginine biosynthesis IV, NAD biosynthesis from 2-amino-3-carboxymuconate semialdehyde and riboflavin, FMN and FAD transformations pathways (NES score > 1.345, *p-value* < 0.05; Supplementary Table 4 and Figure 1B). Interestingly, two main metabolites implicated in these pathways, AMP and FMN are deregulated. Differential metabolite abundance analysis showed that NM1 KO Cells exhibit significantly higher levels of AMP (*t* = 3.07, *p-value* = 0.015), and reduced levels of FMN (*t* = -3.43, *p* value = 0.009; Supplementary Table 2 and 4, and Figure 1D). Elevated AMP and reduced FMN levels can significantly modulate platelet activation through the interplay of cyclic AMP (cAMP) and phosphoinositide signaling pathways. cAMP regulation is crucial, as it inhibits key processes in platelet activation, including calcium mobilization and integrin activation, thereby reducing platelet aggregation and secretion (42, 43). Furthermore, increased cAMP levels suppress G protein-mediated responses, leading to decreased thrombin-induced secretion and aggregation (43). Reduced levels of FMN can disrupt calcium signaling, further influencing platelet activation dynamics (44).

Altogether these observations suggest that loss of NM1 is associated with the differential abundance of metabolites important for hemostasis and, in particular abundance of negative regulators of platelet aggregation.

### NM1 deletion leads to deregulated expression of cytokines and proinflammatory chemokines

Since platelet activation and aggregation play a role in modulating immune responses and inflammation we next studied, if NM1 expression levels affect cytokines and proinflammatory chemokines production using WT, NM1 KO, and rescued NM1 KO MEFs with exogenous NM1 expression (KO+NM1). We measured the amounts of secreted cytokines and chemokines in growing media by using BioLegend LEGENDplex bead-based immunoassays. The results show that most tested cytokines and chemokines levels were altered in the NM1 KO background (Figure 1E and 1F). Except for IL-1β and IL-11, all other cytokines tested including IL-1α, IL-3, IFN-β, IL-12p40, IL-23, IL-7, IL-33, GM-CSF, IL-12p70, IL-27, TSLP were suppressed in the absence of NM1, with the majority being rescued by reintroducing exogenous NM1 in the KO background (KO+NM1) (Figure 1E). Several chemokines show the same profile and their secretion to media is suppressed in KO cells and restored in KO+NM1 cells (RANTES, Eotaxin, MCP-1, MIP-1β, BLC). Only two chemokines show opposite effects in expression and are increased in NM1 knock-out cells (MIP-3α, KC). The chemokines IP-10, MIP-1α and LIX do not show any change upon deletion of NM1 but their expression is increased by the reintroduction of NM1 in the KO background, and chemokines TARC and MDC are suppressed in KO cells but overexpression of NM1 did not lead to restoration of their levels (Figure 1F). These cytokines and chemokines are essential for the proliferation and differentiation of various hematopoietic cells contributing to immune cell development and inflammation processes. Therefore, their broad deregulation in NM1 KO cells suggests the direct involvement of NM1 in the development and functionality of immune system processes. As cytokines and chemokines are secreted by many cell types and serve as broad signaling molecules, we wanted to test whether profiles observed in NM1 knockout cells are specific to NM1 protein or whether their profile changes regardless of the mutated gene. To test this, we used cells lacking beta-actin that exhibit pleiotropic effects on transcription, chromatin remodeling, DNA damage response, and mitochondrial function similar to NM1 protein. We measured cytokines and chemokines produced by these cells and cells and compared them to wild-type control cells. In comparison to NM1 deletion, the deletion of beta-actin has only a marginal effect on cytokine production with only IL-1α being upregulated and GM-CSF being downregulated in beta-actin knock-out cells (Supplementary Figure 1E). In contrast, the expression of several chemokines is changed in beta-actin knockout cells (RANTES, TARC, KC, IP-10, and BLC are suppressed and MIP-3α, MCP-1, MIP-1α, and LIX are upregulated) (Supplementary Figure 1F), however, these changes do not correlate with data observed in NM1 KO cells, supporting a specific role of NM1 in the production of cytokines and chemokines that might act as signaling molecules during hematopoiesis crosslinking platelets activation signaling and immune responses.

### NM1 deletion affects the transcriptomic profile of hematopoietic tissues

To find out if NM1 plays a role in regulating hematopoiesis in vivo, we next isolated RNA from bone marrow, spleen, and peripheral blood from WT and NM1 KO mice for deep sequencing. Principal Component Analysis (PCA) for each tissue shows that there is significant differential gene expression between WT and KO conditions. Moreover, while WT samples show relatively high consistency between the replicates, NMI KO tissue samples show higher variability suggesting that NM1 deletion leads to a general imbalance in global gene expression in the selected tissues (Figure 2A). Due to high variability between NM1 KO replicates, the total number of statistically significant differentially expressed genes is lower in comparison to previously published RNA sequencing data from stable cell lines (7, 8). In bone marrow tissue, there is an equal number of significantly up-regulated (374 genes) and significantly down-regulated genes (358 genes). In the spleen, the majority of differentially expressed genes are downregulated (1134 genes) and only 367 genes are upregulated. On the contrary, in blood, there are 248 upregulated genes and only 52 downregulated genes (Figure 2B). We next compared all differentially expressed genes between the tissues to uncover whether the same set of genes is affected between tissues (Figure 2C). The Venn diagram shows that the majority of differentially expressed genes are specific for each tissue with only a minor part being shared between the groups. However, the shared genes show a high degree of similarity in expression patterns between bone marrow and blood samples (Figure 2D), moderate similarity between bone marrow and spleen samples (Figure 2E), and no similarity between spleen and blood samples (Figure 2F).

**Figure 2.**
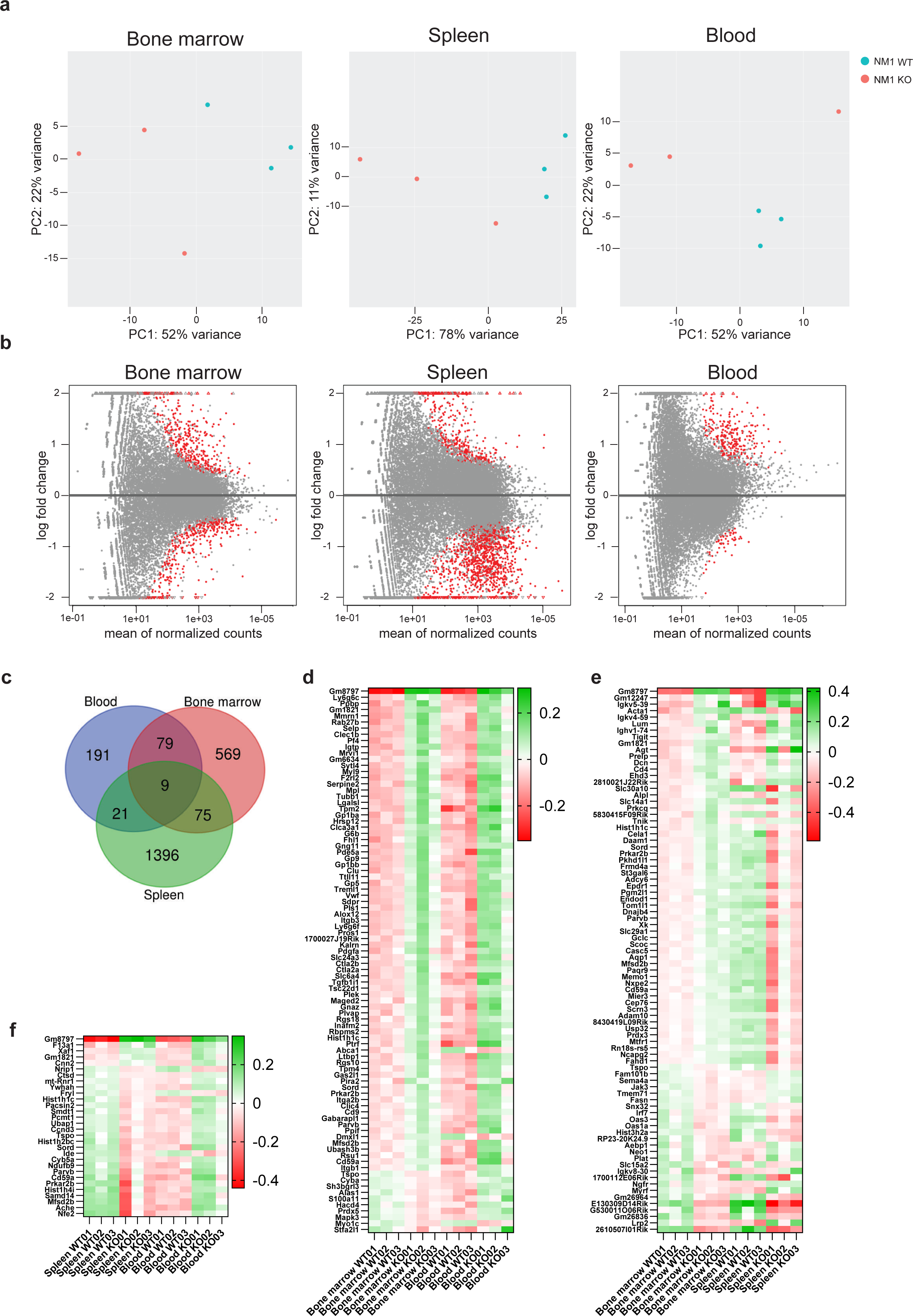
NM1 deletion affects the transcriptomic profile of hematopoietic tissues. **(a)** PCA plot showing the distribution of RNA-seq samples isolated from NM1 WT and KO bone marrow, spleen, and blood samples. **(b)** MA plot visualization of the differences in gene expression between experimental conditions as a function of log fold change versus the mean of normalized counts. Each dot represents a single gene expression profile with only red-marked genes to be significantly differentially expressed. **(c)** Venn diagram showing the intersection in all differentially expressed genes between hematopoietic tissues. **(d)** Expression heatmap of all differentially expressed genes shared between bone marrow and blood samples organized in descending order. **(e)** Expression heatmap of all differentially expressed genes shared between bone marrow and spleen samples organized in descending order. **(f)** Expression heatmap of all differentially expressed genes shared between spleen and blood samples organized in descending order.

This suggests that certain phenotypic changes upon NM1 deletion persist from early hematopoiesis in the bone marrow to differentiated cell types in peripheral blood, while some others are affected by the local environment and/or metabolic requirements of a given cell type.

### NM1 regulates hemostasis and platelet activation in mice

Following RNAseq analysis, we first performed gene ontology analysis of all differentially expressed genes to assess the biological processes and pathways specifically affected by NM1 deletion in bone marrow, where hematopoietic differentiation occurs (Figure 3A). Among the most affected ones, we found biological processes associated with extracellular matrix and cell adhesion, immune system processes, platelet activation and coagulation, osteoclast differentiation, and regulation of PI3K-AKT, Protein kinase B (PKB), MAPK-ERK or Rap1 signaling pathways. To identify the processes activated or suppressed following NM1 deletion, we performed gene ontology analysis with only up-regulated (Figure 3B) or down-regulated genes (Figure 3C). While the majority of aforementioned processes are activated upon NM1 deletion, osteoclast differentiation and processes related to the immune system and immune response are suppressed in NM1 KO mice. This suggests that depending on the process, NM1 may have an antagonistic effect on the expression of specific genes. All affected pathways have been shown to regulate hematopoiesis stem cell maintenance and lineage development. PI3K-AKT signaling is a pro- survival and pro-proliferative pathway that together with PKB signaling regulates the proliferation and renewal of HSCs, and contributes to their differentiation. If deregulated, PI3K-AKT signaling dramatically affects hematopoiesis leading to various hematologic malignancies (45–47). Rap1 signaling pathway regulates extracellular matrix formation and cell adhesion and it is important for the differentiation of megakaryocytes and platelets via MAPK-ERK pathways (48–50), which is in agreement with observed gene ontology terms found in the upregulated genes related to the extracellular matrix, cell adhesion, and platelet activation (Figure 3B).

**Figure 3.**
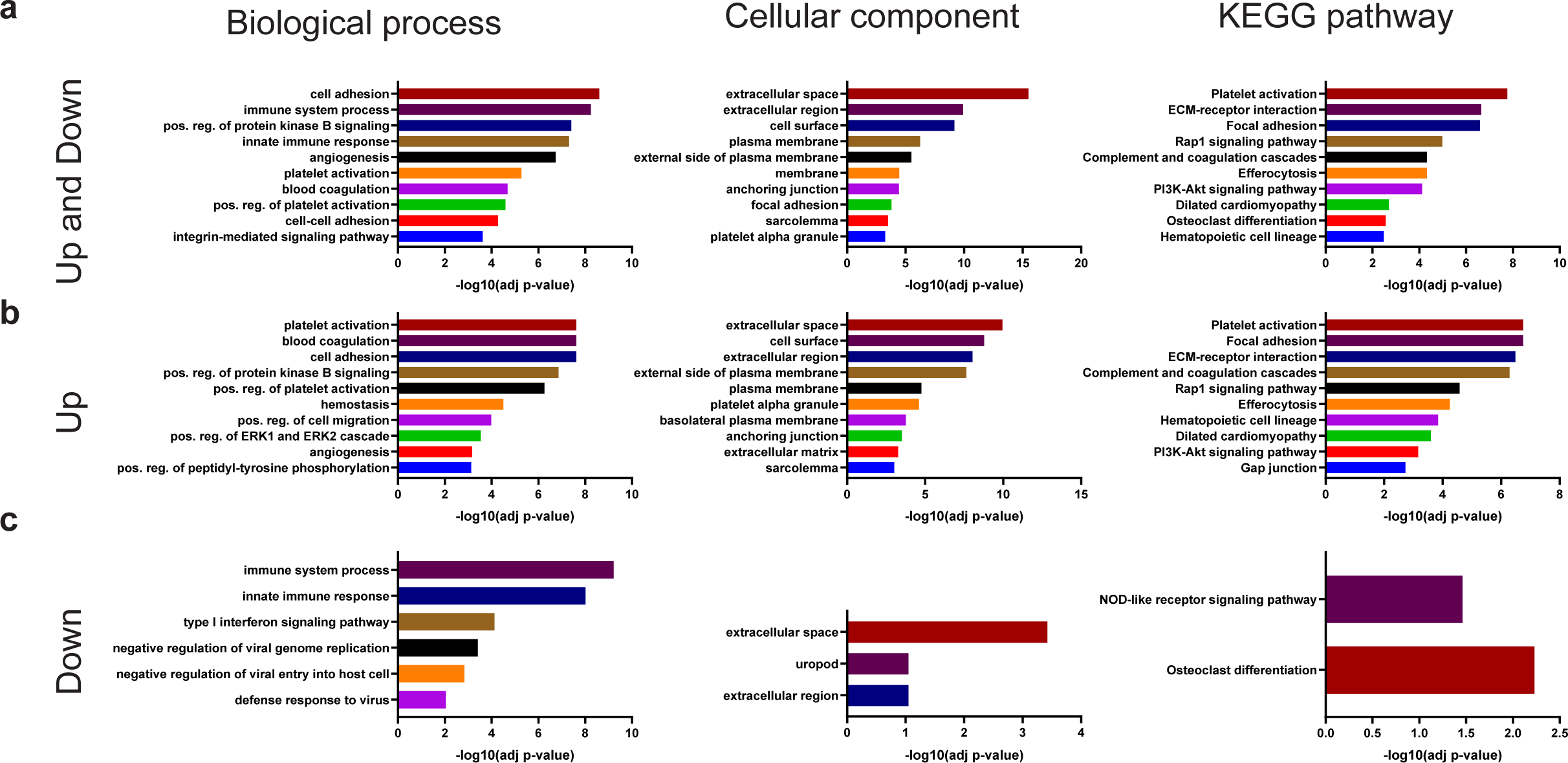
Gene ontology analysis of bone marrow samples isolated from NM1 WT and KO mice. **(a)** Gene ontology analysis based on all differentially expressed genes between experimental conditions shows biological process-, cellular component-, and KEGG pathway-associated gene ontology terms plotted in descending order based on their significance. **(b)** Gene ontology analysis based on only upregulated genes in NM1 KO samples. **(c)** Gene ontology analysis based on only downregulated genes in NM1 KO samples.

Next, we focused on differentially expressed genes associated with hemopoietic differentiation, platelet activation, and blood coagulation and performed STRING analysis of these genes to produce a gene interaction map based on text mixing, experimental data, co-expression, and database search (36) (Figure 4A). This analysis led to the discovery of two main clusters – genes directly involved in platelet activation and blood coagulation processes (Cluster I), and genes involved in cell signaling cascades (Cluster II). The first group contains collagen genes (Col1a1, Col1a2), which are recognized by von Willbrand factor (Vwf), and genes expressing platelet glycoproteins (GP5, GP9, Gp1ba, GP1bb) which attach platelets to collagen network directly or via Vwf. Integrins (Itgb3, Itga2b, Itgb1, Itga6) present on the platelet plasma membrane then mediate platelet aggregation via binding to extracellular proteins such as fibrinogen to form blood clots (51). Several other factors regulate platelet activation, including platelet factor 4 (Pf4) released during platelet aggregation to neutralize the anticoagulant effect of heparin, Multimerin 1 (Mmrn1) which stabilizes factor V in platelets, and the CD9 antigen (Cd9) that controls platelet activation and aggregation via cell adhesion and others (Serpine2, Serping1, Thbd, Tfpi, G6b, F2Rl3, F2Rl2, Pros1). The second group of genes identified in the STRING analysis is related to different cellular signaling cascades having a role during cell adhesion, growth, and proliferation. This group contains genes expressing Tyrosine-protein kinases (Mapk13, Mapk3, Src, Mertk), thyrosine-protein phosphatases (Ptprj), receptors and ligands for thyrosine kinases (Kitl, Axl, Gas6), and G-protein coupled signaling proteins (Adyc6, Adyc5, Il1A, Gnas, P2Ry1, Gucy1a3, Gucy1b3). The expression profile of all differentially expressed genes across experimental samples associated with GO terms “Platelet activation”, “Hematopoietic cell lineage”, and “Blood coagulation” is shown as a heatmap in descending order (Figure 4B).

**Figure 4.**
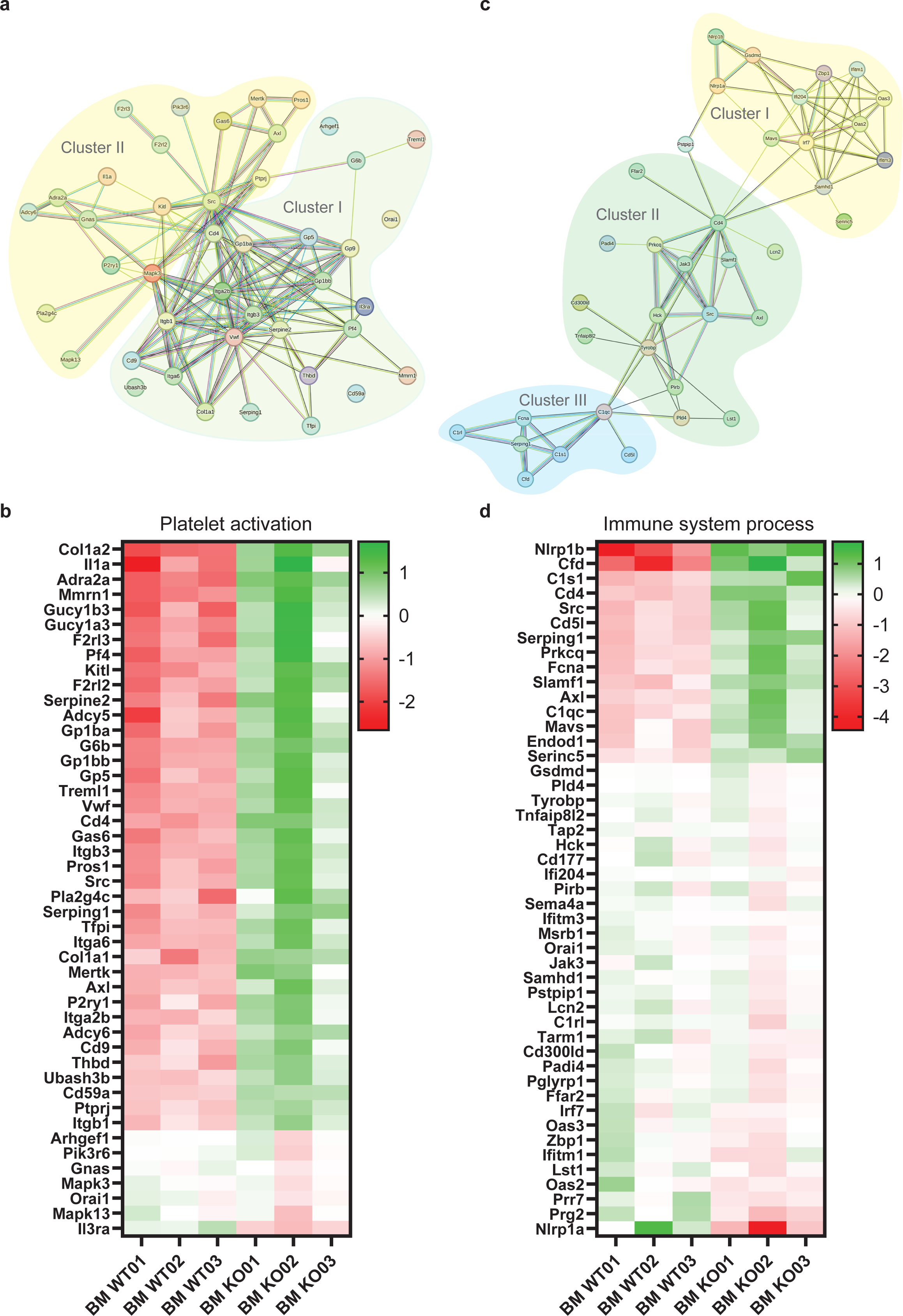
Gene expression analysis of genes associated with platelet activation and immune system process in the bone marrow. **(a)** String analysis diagram of all genes associated with a KEGG GO term “Platelet activation”, a Biological process GO term “Platelet activation”, KEGG GO term “Haemopoietic cell lineage”, and Biological GO term “Blood coagulation”. **(b)** Heatmap of all differentially expressed genes between NM1 WT and KO bone marrow samples associated with Platelet activation GO terms used in String analysis. **(c)** String analysis diagram of all genes associated with GO term “Innate immune system”. **(d)** Heatmap of all differentially expressed genes between NM1 WT and KO bone marrow samples associated with Innate immune system GO term used for the String analysis.

Since platelet activation is a key component of the complex response to vascular injury in peripheral tissues, we next performed transcriptional profiling of WT and NM1 KO peripheral blood, the site of action for circulating blood cells. We found a relatively low number of differentially expressed genes (∼300) with the majority of them being upregulated. Similarly to bone marrow, gene ontology analysis shows the majority of upregulated GO terms are associated with blood coagulation, platelet activation, hemostasis, and related processes such as rearrangement of the actin cytoskeleton, cell-matrix adhesion, and integrin-mediated signaling of all differentially expressed genes (Supplementary Figure 2A and 2B). We next plotted all genes associated with GO terms “Blood coagulation” and “Platelet activation” as heatmaps in descending order (Supplementary Figure 2C and 2D respectively). We found several platelet activation and coagulation genes (Vwf, F2rl2, Serpine2, Pros1, G6b), platelet glycoproteins (Gp1bb, Gp5, Gp9, Gp1ba, Gp1bb), integrins (Itga2b, Itgb3, Itgb1) to be upregulated in NM1 KO blood samples as well as in bone marrow. Above that, other platelet glycoproteins (Gp6), coagulation factors (F5, F10, F13a), platelet activation and adhesion genes (Ptgs1, Fermt3, Anxa5), signaling proteins (Rasgrp2, Rap1b, Rasgrp1), cytoskeletal proteins (Actb, Actg1, Mylk) and others to be specifically changed in NM1 KO blood samples (Supplementary Figure 2C and 2D).

Next, we assessed the impact of NM1 deficiency on primary hemostasis, by measuring bleeding time in WT and NM1 KO mice (Figure 5A). Surprisingly, even though NM1 deletion leads to hyperactivation of platelet activation cascade, NM1 KO mice show significantly prolonged bleeding time in comparison to WT (p = 0.0326), indicating a general impairment of clotting efficiency in the absence of NM1. The mean bleeding time was 225.0 ± 35.18 sec in WT mice, whereas NM1 KO mice exhibited a significantly increased mean bleeding time of 322.5 ± 101.1 sec (Figure 5B and Supplementary Figure 3). To investigate whether in the NM1 KO condition platelet homeostasis is also affected, we next performed a complete blood count (CBC) analysis in WT and KO mice. Given the prolonged bleeding time observed in KO mice, we aimed to determine whether altered platelet numbers could contribute to impaired hemostasis. CBC analysis revealed a significant reduction in platelet counts in KO mice compared to WT controls (p = 0.05). The mean platelet count in WT mice was 174.7 ± 48.79 ×10³/µL, whereas KO mice exhibited a markedly lower mean platelet count of 52.25 ± 26.85 ×10³/µL (Figure 5B). In contrast to white blood cell (WBC) counts which were not significantly different between the groups (Figure 5B and Supplementary Figure 3), CBC analysis also revealed a significant drop in red blood cell (RBC) count (Figure 5B), a decrease in hematocrit (HCT) levels and an increase in the average size of red blood cells (MCV) (see Supplementary figure 3) as previously published (29).

**Figure 5.**
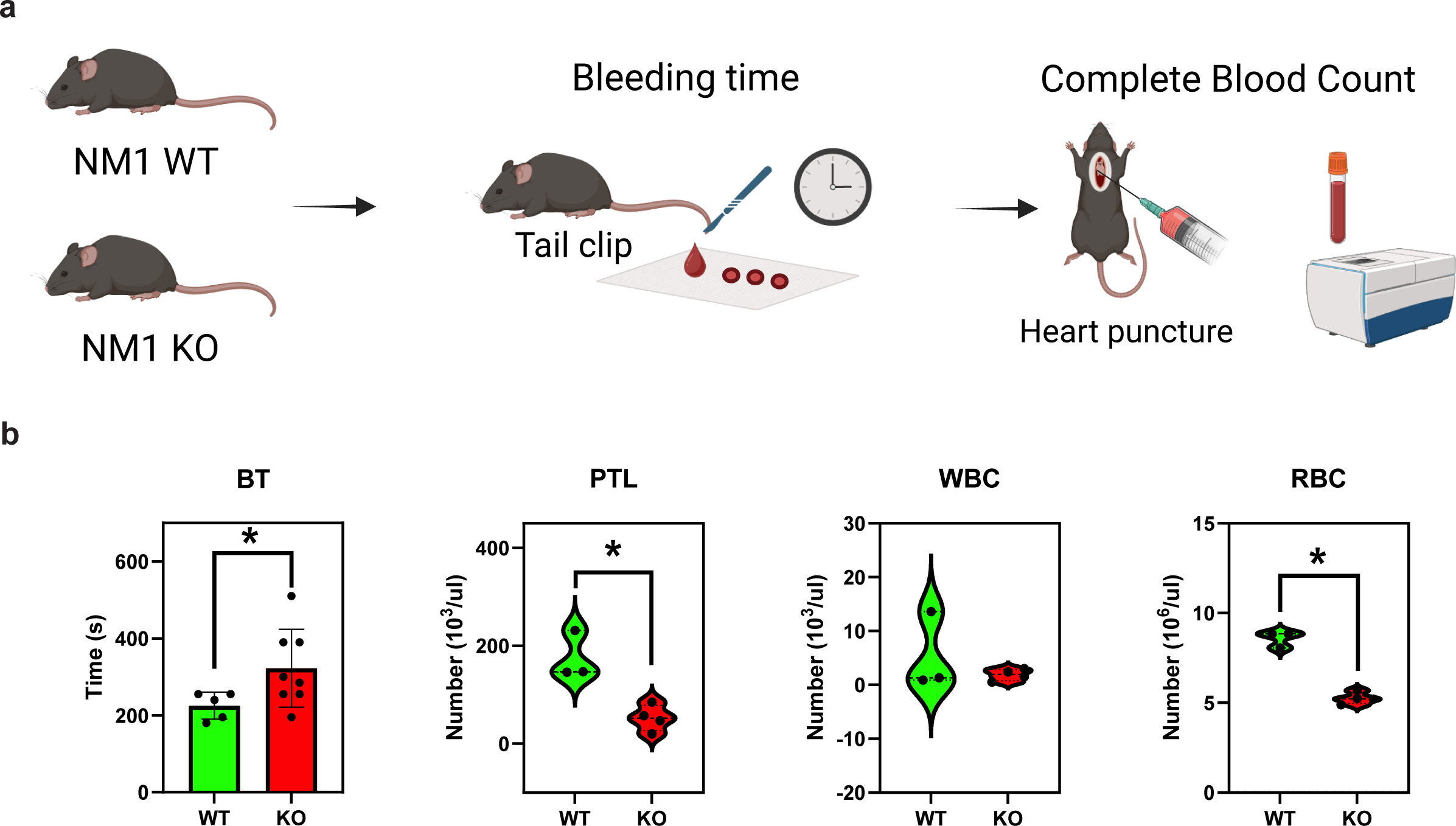
NM1 deficiency leads to prolonged bleeding time and decreased number of platelets. **(a)** Schematic diagram illustrating the experimental procedure, where mice undergo tail cut for bleeding time (BT) measurement. Following this, whole blood is collected via cardiac puncture for subsequent CBC analysis. **(b)** Bar graph (Mean±SD) showing bleeding time (BT) between WT and NM1 KO mice (n = 5 WT, 9 KO). Additional violin plots show PTL, WBC, and RBC for both groups (n = 3 WT, 4 KO). The two-tailed Mann-Whitney U test was used for statistical analysis. * P≤0.05, ** P≤0.001, *** P≤0.001, **** P≤0.0001.

Taken together, the observations from the metabolomics analysis of primary cells, RNA sequencing of hematopoietic tissues, and functional assays point towards the complex role of NM1 in the process of hemostasis with major effect on platelet differentiation and activation.

### NM1 regulates innate and adaptive immune cell differentiation and activation

Following the results from cytokine and chemokine measurements in model cell lines and the abundance of GO terms associated with the immune system in bone marrow, we next performed a String analysis of genes related to the gene ontology term “Immune response process” (Figure 4C). We found that they cluster into three distinguished groups. The first group of genes (Cluster I) is associated with the innate immune system and defense response to viruses correlating with GO terms found in the gene ontology analysis of downregulated genes in NM1 KO mice. Among them, we found the interferon regulatory factor 7 (Ifn7), a key transcriptional regulator of type I interferon-dependent immune responses against DNA and RNA viruses. Compatible with the identification of Ifn7, in the same group we found the mitochondrial antiviral signaling protein (Mavs), a central factor in signal transduction upon viral infection, required for activation of transcription factors regulating interferon beta expression (52). Ifitm1 and Iftm3, interferon-induced transmembrane antiviral proteins which inhibit the entry of the viruses to the host cell cytoplasm, also cluster in the same group, similarly to Serine incorporator 5 (Serinc5), required to restrict incorporation of viral particles into cells. We also found interferon-induced genes such as Oas2 and Oas3, coding for 2’-5’ oligoadenylatesynthetases 2 and 3, which are dsRNA-activated antiviral enzymes that degrade viral RNA (53) and Samhd1 a cellular nuclease with dNTPase activity that reduces cellular dNTP levels to protect cells from retroviral reverse transcription during viral replication as well as Z-DNA-binding protein 1 (Zbp1), a cytoplasmic DNA sensor that detects DNA of viral or bacterial origin. Finally, we identified Nlprp1a and Nlprp1b, two factors that play a crucial role in the formation of the inflammasome in response to pathogens and other damage-associated signals.

In the second cluster (Cluster II) are proteins involved in intracellular signaling cascades and immune regulatory proteins, some of which were already described in platelet activation (Src, Axl, Cd4). In addition to those, we found hematopoietic tyrosine-protein kinase (Hck) which regulates phagocytosis and neutrophil, monocyte, macrophage, and mast cell function during innate immune response. Protein kinase C theta (Prkcq) regulates T-cell activation, proliferation, differentiation and survival. Tyrosine protein-kinase Jak3 mediates essential signaling events in both innate and adaptive immunity and plays a crucial role during T-cell development. The other immune system receptors and regulators found in these clusters are TYRO protein tyrosine kinase-binding protein (Tyrobp), Tumor necrosis factor alpha-induced protein 8-like protein 2 (Tnfaip8l2), leukocyte immunoglobulin-like receptor subfamily B member 3 (Pirb), CMRF35-like molecule 5 (Cd300Id) and leukocyte specific transcript 1 (Lst1).

The third cluster of genes derived from the STRING analysis relates to the immune complement system to enhance the phagocytosis and clearance of microbes and damaged cells. It contains complement factors Cfd, C1s1, C1qc, C1rl and regulatory proteins Serping 1, which regulates the activation of the complement system by inhibiting C1 factor (54) and Ficolin-1 (Fcna), which is extracellular lectin functioning as a recognition receptor in innate immunity in the lectin complement system pathway (55). Expression profile across biological replicates for all significantly different genes classified under GO term “Immune system process” is shown as a heatmap in descending order (Figure 4D). As suggested by the gene ontology analysis, the majority of genes related to the immune process is downregulated in NM1 KO cells but the differences between the genotypes are less prominent in comparison to differential gene expression related to platelet activation (Figure 4D).

Since many immune cells reside and get activated in secondary lymphoid organs, we next studied potential changes in immunity pathways in spleen tissue isolated from NM1 WT and KO mice. Results from gene ontology analysis show that the most significantly dysregulated biological processes and pathways are associated with the cell cycle and cell division regulation, DNA replication, and chromosome segregation, as well as immunoglobulin production and immunoglobulin-mediated immune response (Figure 6A). To define which biological processes and pathways are positively or negatively affected by NM1 deletion, we performed separate gene ontology analyses of only upregulated and only downregulated genes in the NM1 KO spleen respectively. In contrast to bone marrow tissue, immune system-related genes are predominantly upregulated in the NM1 KO spleen tissue (Figure 6B), and downregulated genes are mostly associated with the cell cycle and cell division GO terms (Figure 6C). To understand the differences in immune-related processes between the bone marrow and the spleen, we took all differentially expressed genes listed under GO terms “immunoglobulin mediated immune response”, “immunoglobulin production”, “immune response”, “immune response process”, “adaptive immune response”, “T-cell receptor signaling pathways”, and “T cell activation” and performed the STRING analysis of protein interactions. The majority of differentially expressed genes that could not be mapped during the analysis are variable or constant gene segments of immunoglobulin heavy and light chains that do not possess any particular function per se (Figure 7A). They are randomly combined by V(D)J somatic recombination into functional heavy and light chains to produce variable antibodies. V(D)J recombination is executed already during lymphoid development of B cells in bone marrow but antibodies are presented only in mature naïve B cells present in the spleen and other tissues. Therefore, even though this particular immunoglobulin gene expression profile is found in the spleen tissue, the phenotype itself originates in the bone marrow. So, we looked at the expression profile of V(D)J recombination-specific genes in bone marrow (Supplementary Table 5), and found Rag1 and Rag2 genes to be significantly upregulated. These recombination-activating genes 1 and 2 initiate V(D)J recombination by introducing double-strand breaks between the recombination signal sequence (RSS) and a coding segment of a particular immunoglobulin heavy or light chain gene (56) and their overexpression leads to lymphocyte malignancies and immunodeficiency (57–59). However, it is not known whether there is a direct link between increased Rag1 and Rag 2 expression in the bone marrow and a particular combination of the immunoglobulin heavy and light variable gene segments found in the NM1 KO spleen.

**Figure 6.**
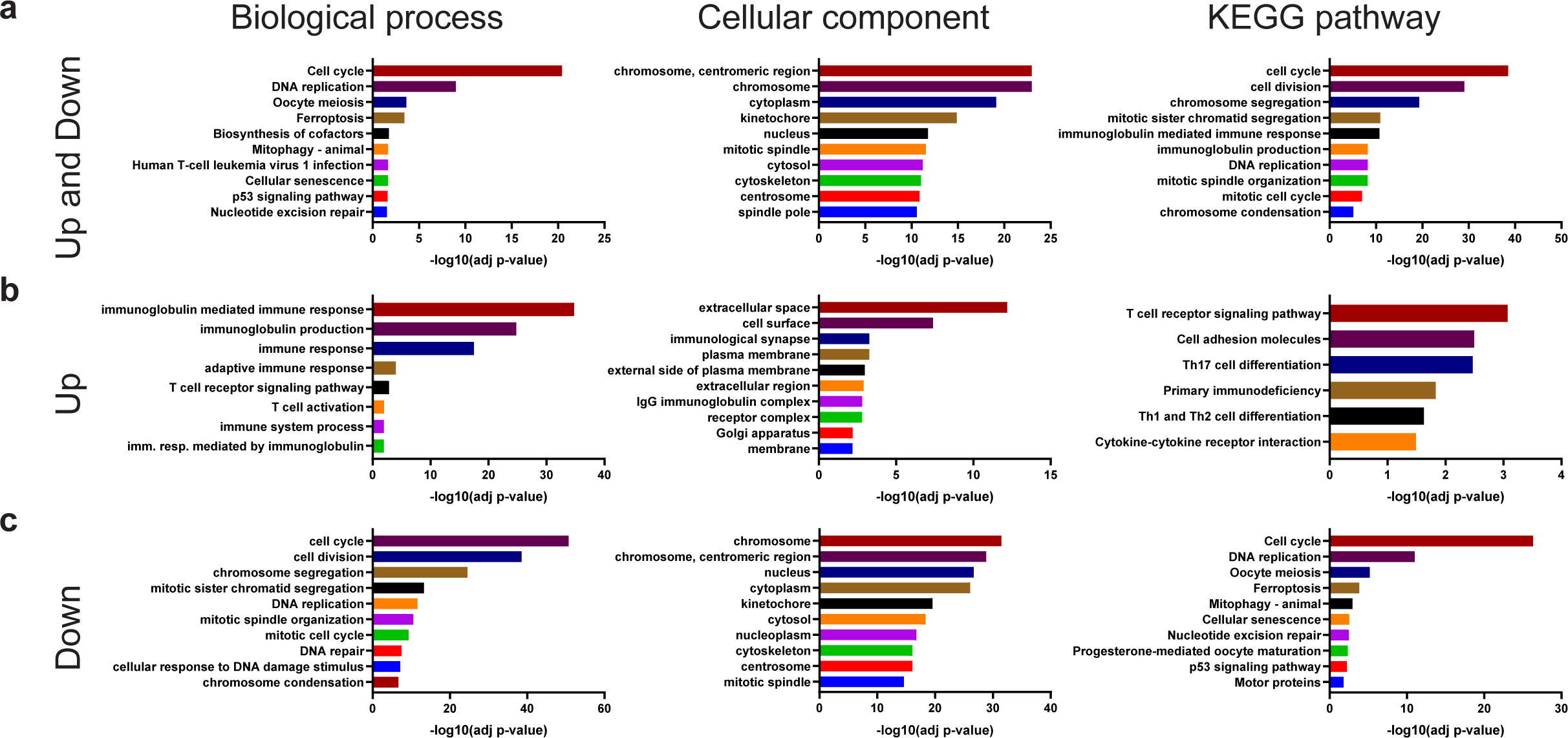
Gene ontology analysis of spleen isolated from NM1 WT and KO mice. **(a)** Gene ontology analysis based on all differentially expressed genes between experimental conditions shows biological process-, cellular component-, and KEGG pathway-associated gene ontology terms plotted in descending order based on their significance. **(b)** Gene ontology analysis based on only upregulated genes in NM1 KO samples. **(c)** Gene ontology analysis based on only downregulated genes in NM1 KO samples.

**Figure 7.**
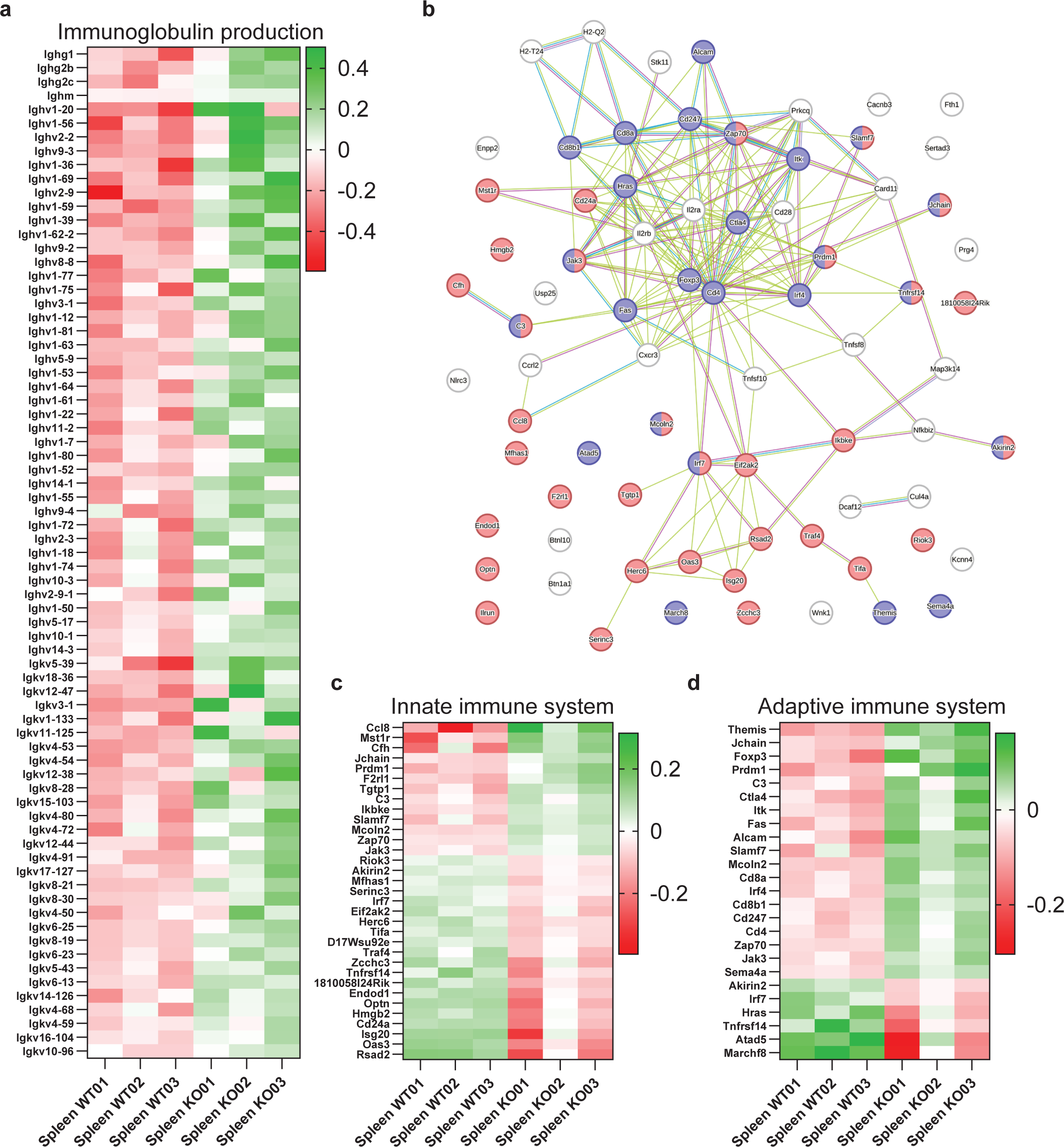
Gene expression analysis of genes associated with the innate and adaptive immune system in the spleen. **(a)** Heatmap of all differentially expressed genes between NM1 WT and KO spleen samples associated with Immunoglobulin production which could not be mapped in String analysis **(b)** String analysis diagram of all genes associated with GO term “Immune system”. Genes marker red are associated with the innate immune system while genes marked blue are associated with adaptive immune system. **(c)** Heatmap of all differentially expressed genes between NM1 WT and KO spleen samples associated with the innate immune system. **(d)** Heatmap of all differentially expressed genes between NM1 WT and KO spleen samples associated with the adaptive immune system.

Next, we plotted all other immune-related genes into an interaction map based on the experimental data, text mining, co-expression, and database search and divided them by their role in innate (red) or adaptive (blue) immune systems (Figure 7B). The proteins marked by both colors have a dual role in both types of immunity and proteins marked white are additional factors present in immune cells without a direct link to either innate or adaptive immune systems. The only visible cluster of innate immune genes is localized around Interferon regulatory factor 7 (Irf7) protein which is the master regulator of type I-interferon genes and interferon-stimulated genes. The other proteins in the cluster are different interferon-stimulated enzymes important for the innate immune response against viruses and bacteria (Eif2ak2, Herc6, Oas3, Rsad2, Isg20, Serinc), and proteins activating proinflammatory NF-kappa-

B signaling (Ikbke, Traf4, Tifa). Other innate immune system-related genes are different regulatory and signaling proteins (Riok3, Mfhas1, Ilrun, Zcchc3, Ccl8, Mst1r, F2rl1, Slamf7, Mcoln2), developmental and differentiation genes (Akirin2, Tnfrsf14, Cd24a, Prdm1, Zap70, Jak3), and additional relevant genes.

Adaptive immune system-related genes (blue) form a big interconnected cluster of genes regulating different aspects of T-cell and B-cell biology. To differentiate between them, we marked all genes associated with B cells and T cells separately based on the 5 most significant GO terms associated with each cell type (Supplementary figure 4A and 4B respectively) and found that the majority of differentially expressed genes are associated with the T cell development and specification (Themis, Foxp3, Prdm1, Fas, Jak3, Sema4a), cell surface receptors (Cd4, Cd8a, Cd8b1, Cd247) and T cell activation and signaling (Itk, C3, Ctla4, Alcam, Slamf7, Irf4, Zap70, Tnfsrf14).

We then plotted all Innate immune system-related and adaptive immune system-related genes into a heatmap in descending order based on their expression profile in the NM1 KO spleen (Figure 7C and 7D). Interestingly, the majority of innate immune system genes are suppressed in the NM1 KO spleen similarly to bone marrow, with some genes being shared between both tissues (Irf7, Endod1, Oas3) (Figure 7C). Many upregulated genes have a dual role in both types of immune response and are therefore present also in the adaptive immune system heatmap. In contrast to the innate immune system, adaptive immune system genes are mostly upregulated suggesting some antagonistic effect of NM1 on differentiation and activation of both immune systems in mice. Interestingly, based on the expression profile it seems that there is an overrepresentation of several types of regulatory T cells (Tregs). While Cd4 and Cd8 are general markers of mature T cells, in combination with other factors define specific T-cell subpopulations. Upregulated Foxp3 and Il2ra (CD25) are markers of Cd4+ Foxp3+ CD25+ (60), and Cd8+ Foxp3+ CD25+ Treg cells (61). Upregulated Il2rb (CD122) is a marker of Cd8+ Cd122+ Treg cells which are further specified by increased expression of Pdcd1 (Pd1) which is also upregulated (but not statistically significant) in NM1 KO spleen (62–64) (Supplementary table 5). Ctla4 is upregulated in activated T-cells but is also constitutively expressed in Cd8+ Ctla4+ Treg cells (65, 66) (Figure 7D).

We conclude that NM1 plays an important, possibly antagonistic role in innate and adaptive immune systems. Given its effect on platelet differentiation and activation, we propose that NM1 plays a general role in the communication and coordination of immune and hemostatic processes in mice.

## Discussion

After the cell nucleus, mitochondria are probably the second most studied cellular organelle due to their role as a powerhouse producing energy via oxidative phosphorylation in healthy cells. In contrast to oxidative phosphorylation, aerobic glycolysis has long been considered a metabolic pathway typical of cancer cells or a consequence of aberrant mitochondrial function. However, discoveries over the last decade have shown that aerobic glycolysis is a full-featured pathway that does not only produce energy but also provides specific byproducts and signaling molecules or specific intracellular and extracellular conditions to many cell types. Moreover, there is a clear interplay between oxidative phosphorylation and aerobic glycolysis and the energy production pathways can change even in the same cell type based on the actual needs, cellular activity, or differentiation status. The regulation of this metabolic switch is still poorly understood but it is getting gradual attention as it is associated with many metabolic disorders. Recently, we discovered that NM1 plays a critical role in this process and its mutation or deletion leads to a metabolic switch from oxidative phosphorylation to glycolysis, and subsequent tumorigenesis in mice (8). Because of these observations, we hypothesized that NM1 may be a general factor regulating the switch between oxidative phosphorylation and glycolysis in multiple tissues during the differentiation or activation of specific cell types.

The bone marrow is a very heterogeneous tissue giving rise to several, mostly hematopoietic cell types with very specific functions and metabolic needs, and thus it is an interesting non-cancerous model to study whether NM1 depletion affects cell differentiation possibly by affecting the interplay between oxidative phosphorylation and glycolysis. Several studies have suggested the involvement of both actin and myosin in the regulation of the immune system (67). In addition, initial phenotyping of NM1 KO mice as well as this study, revealed a decreased total number of red blood cells with higher corpuscular volume and hemoglobin content and decreased bone mineral density suggesting that NM1 can play a role in the differentiation of cells in bone marrow and other hematopoietic tissues (29). Therefore, we performed transcriptomic analysis of the bone marrow, spleen, and peripheral blood isolated from NM1 WT and KO mice to discover gene programs affected by NM1 deletion. Regarding decreased erythrocyte numbers observed in NM1 KO mice, gene programs involved in erythrocyte differentiation are not significantly affected. Although we have limited mechanistic insights at this stage, this result suggests that NM1 may not have a specific role in erythrocyte differentiation or there are compensatory mechanisms upon NM1 deletion. Such mechanisms are observed in the so-called “sports anemia” when trained athletes have decreased hematocrit but the total mass of red blood cells and hemoglobin levels are higher (68). The lack of a dominant effect on transcription profiles during erythropoiesis could also be due to the very metabolic setup of differentiated erythrocytes. Even though there is a short stage during differentiation which depends on oxidative phosphorylation, final red blood cells fully depend on glycolysis (11). As NM1 deletion pushes cells toward glycolytic metabolism, erythrocytes should not be affected as much since mature cells do not depend on oxidative phosphorylation. Similarly, osteoclasts which normally use oxidative phosphorylation as the primary energy source and glycolysis only upon activation to increase local acidosis (20), are transcriptionally suppressed in NM1 KO mice but NM1 KO mice have decreased bone mineral density (29). Therefore, while differentiation of osteoclasts can be negatively affected by NM1 loss, glycolysis-dependent bone resorption could be elevated in NM1 KO mice as seen in osteoporotic patients with hyperactive osteoclasts with increased glycolysis and higher extracellular acidity needed for bone resorption (20, 69).

The most affected pathways in the tested tissues were related to platelet activation and innate and adaptive immune systems. Interestingly, even though the blood cell count shows a significantly lower number of platelets in circulating blood and increased bleeding time in NM1 KO mice, the platelet activation cascade was upregulated in the bone marrow and peripheral blood. Under normal conditions, platelets are formed from proplatelet protrusions of megakaryocytes in the bone marrow. Upon their release into the bloodstream, they are activated at the place of vessel wall injury leading to the reorganization of their morphology and the formation of platelet plugs (70). Therefore, their decreased number and hyperactivation already in the bone marrow suggest general deregulation of platelet biogenesis in NM1 KO mice. We hypothesize that this is due to different metabolic requirements during platelet differentiation and activation. Megakaryocytes depend on both glycolysis and OXPHOS for energy, with glycolysis supporting early differentiation and polyploidization and later proplatelet formation when ATP is needed for cytoskeletal remodeling and formation of the membrane protrusion for platelet production (71). However, mitochondria-dependent reactive oxygen species production is crucial for the initiation of thrombopoiesis, suggesting that functional OXPHOS plays a role in proplatelet formation (72). Subsequently, circulating platelets depend again on both types of metabolism, but glycolysis is required for their activation. Therefore, NM1 loss-dependent decrease in mitochondrial OXPHOS metabolism could lead to decreased proplatelet formation in megakaryocytes and subsequent reduction of circulating platelets, with glycolytic metabolism pushing them towards activation regardless of the external stimuli. Proplatelet formation, followed by platelet segregation and platelet activation, depends on the rearrangements of the actin cytoskeleton and plasma membrane. As NM1 directly links the plasma membrane to the underlying cytoskeleton (4), phenotypes observed in platelets may arise not only due to deregulated metabolism but also due faulty plasma membrane dynamics caused by NM1 loss. Above the main role in blood coagulation, there is also increasing evidence of platelet communication with immune cells in inflammation processes, osteoclast differentiation, and cancer development and progression (73–76). This could be directly linked to phenotypes observed in innate and adaptive immune responses. As mentioned before while innate immune system is suppressed in NM1 KO mice, adaptive immune system and expression of specific sets of immunoglobulins are upregulated. While this can be explained by metabolic reprogramming of given cells and a need for oxidative phosphorylation for differentiation of innate immune monocytes or dendritic cells and glycolysis needed for activation of B and T cells (12, 13). However, it is also plausible that NM1 affects the destiny of these cells differently. Cell metabolism and, in general, proliferation are regulated by the PI3K/Akt/mTOR signaling pathway, with mTOR activating the expression of many downstream targets. In particular, proliferation is heavily regulated by this pathway, with mTOR activating the expression of many downstream targets, including metabolic gene programs. We found that NM1 is a downstream target of mTOR to regulate the expression of mitochondrial transcription factors, but it also regulates mTOR expression itself in a feedback loop. Upon stimulation, mTOR activates NM1, which binds to the mTOR promoter and induces its expression as long as the stimulus persists. Thus, deletion of NM1 may lead to suppression of mTOR signaling, while another pathway may influence the processes observed in NM1 knockout mice. For example, a delicate balance of the complex system of nutrient sensing and mTOR (mTORC1) signaling is crucial to ensure the appropriate development of dendritic cells (77) The differentiation and survival of dendritic cells rely on the mTOR complex 1 (mTORC1) activation and subsequent expression of the mTORC1 downstream target peroxisomal proliferator-activated receptor γ (PPARγ) which affects cell maturation and function largely through control of lipid metabolism and its function is abrogated by rapamycin, an mTOR/mTORC1 inhibitor (78–80). mTOR was also shown to have a role during the differentiation of B cells and VDJ recombination. Here, the mutations in mTORC complex or treatment with rapamycin have positive effects, and lead to enhanced pro-B cell survival, an increase in RAG1 and RAG2 recombinase expression, and augmented V(D)J recombinase activity (81). As the same phenotypes can be observed in NM1 KO immune cells, NM1 may regulate various aspects of hematopoiesis differentiation as a part of PI3K/AKT/mTOR signaling via direct transcriptional regulation of their metabolism, or more broadly by affecting some mTOR-dependent pathways.

Taken together, we showed that NM1 contributes to global body homeostasis by regulating the differentiation and activity of specific hematopoietic cell types. We hypothesize that NM1 affects the destiny of these cells by regulating their metabolism status, but the direct mechanism remains to be elucidated and will be the objective of future studies.

## Acknowledgments

This work was supported by grants from New York University Abu Dhabi, by Tamkeen under the NYU Abu Dhabi Research Institute Award to the NYUAD Center for Genomics and Systems Biology (ADHPG-CGSB), and by the Sheikh Hamdan Bin Rashid Al Maktoum Award for Medical Sciences to PP. We also acknowledge the institutional support from the Institute of Molecular Genetics of the Czech Academy of Sciences (RVO: 68378050) and the Research Program Strategy AV21 Future of Assisted Reproduction (ART) (AV21-VP38/2025) provided by the Czech Agency of Sciences to TV and PH. RNA sequencing was performed by the NYUAD CGSB Core at New York University Abu Dhabi and Bioinformatics assistance was provided by the NYUAD Bioinformatics Core at New York University Abu Dhabi. We would like to thank Marc Arnoux for assistance with deep sequencing and Nizar Drou for assistance with the analysis.

## Author Contributions

TV designed and performed the majority of experiments, analyzed the sequencing data, and wrote the manuscript together with PP. SK, VF, RS, performed functional assays in mice. WA and YI analyzed the metabolomic data. MEG and JCMT performed cytokine assays. PH provided the NM1 KO mice and contributed to manuscript writing. PP supervised the research project, wrote the manuscript, and analyzed the data.

