## Supplemental material for "Nuclear Myosin 1 regulates platelet activation and immune response: A novel therapeutic target for hematologic disorders"

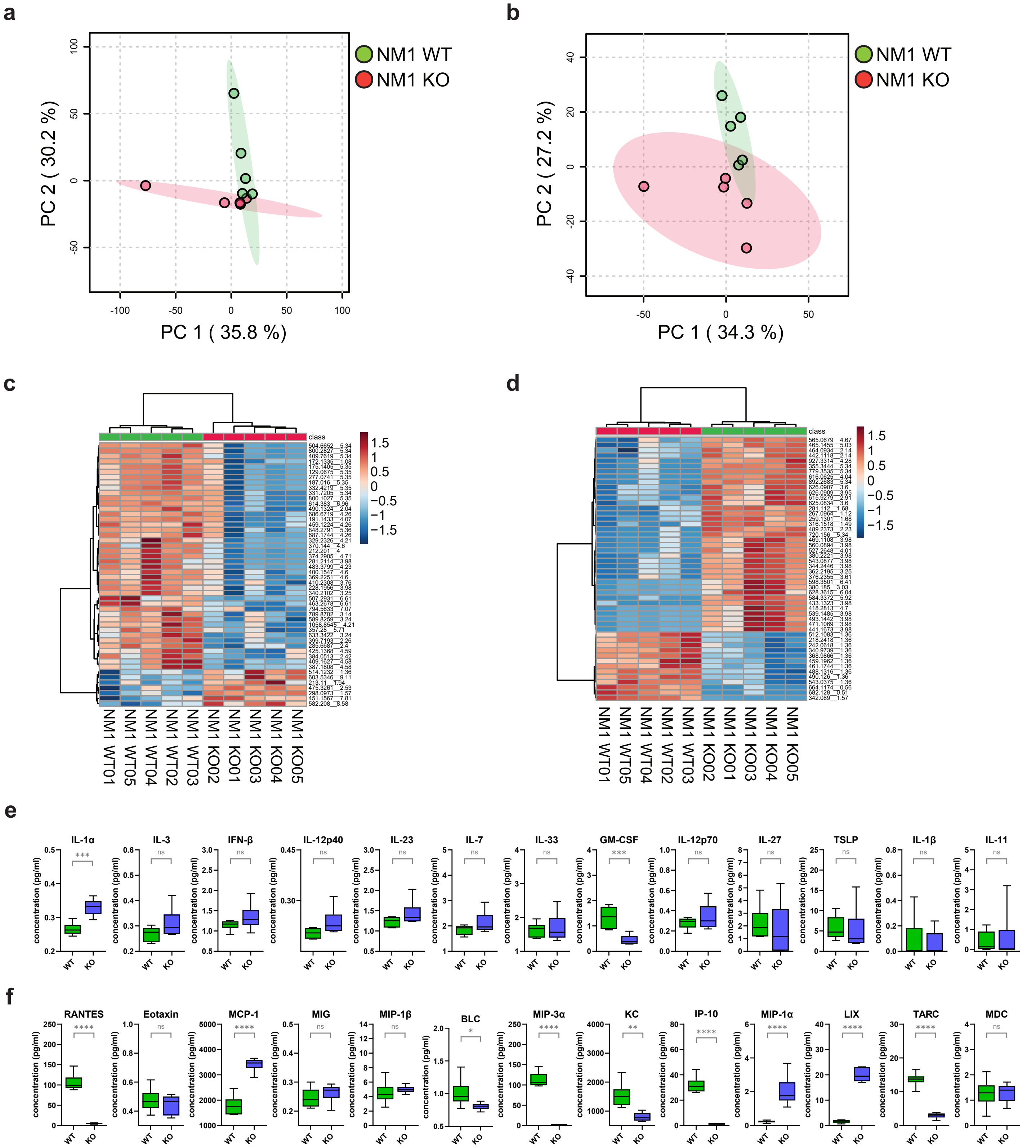


Supplementary Figure 1. Metabolomic profiling and cytokine and chemokine measurement. (a) Principal-component analysis (PCA) of the normalized peak intensity data detected using positive ionization. (b) Principal-component analysis (PCA) of the normalized peak intensity data detected using negative ionization. (c) Clustering of the top 50 significantly differentially abundant metabolic features between the two conditions detected by positive ionization. (d) Clustering of the top 50 significantly differentially abundant metabolic features between the two conditions detected by negative ionization. (e) List of analyzed cytokines in Actin WT and Actin KO cells. ns P > 0.05, *** P≤0.001. (f) List of analyzed chemokines in Actin WT and Actin KO cells. ns P > 0.05, * P≤0.05, ** P≤0.001, **** P≤0.0001.

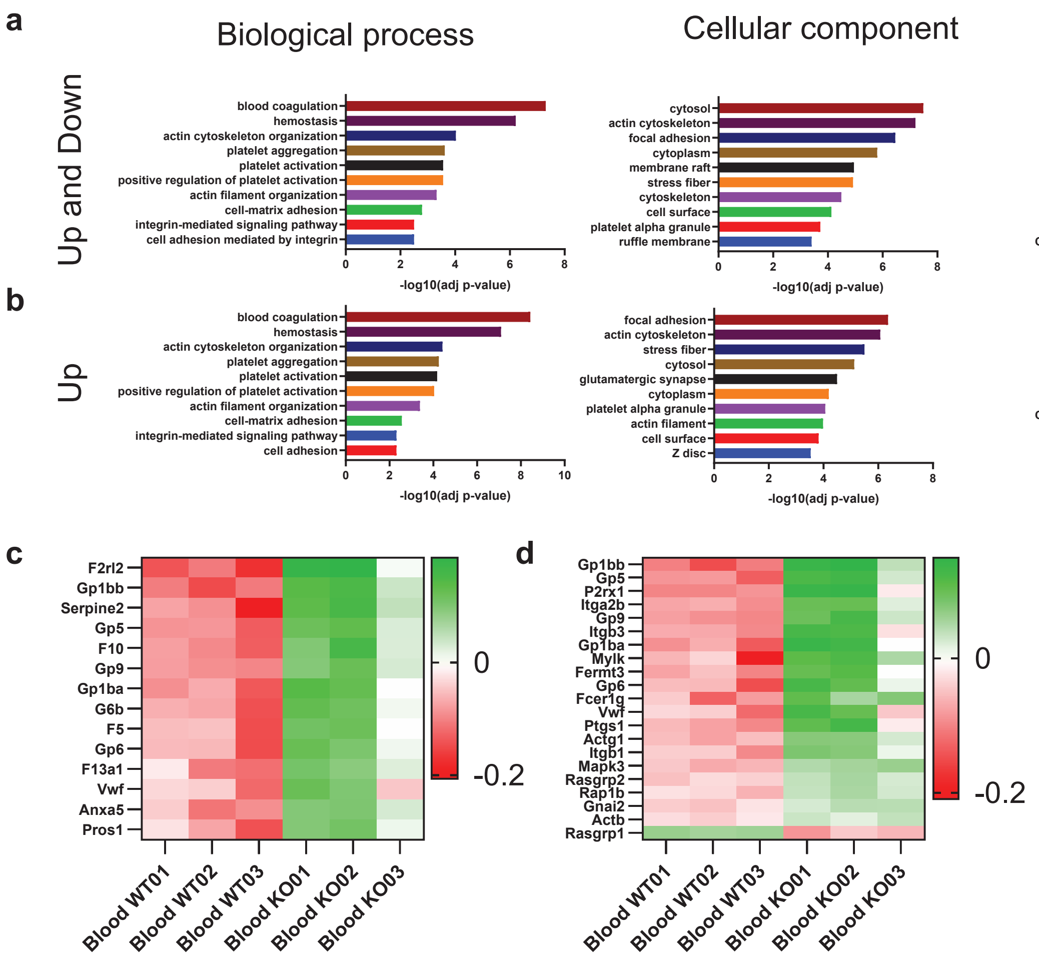


Supplementary Figure 2. GO analysis and gene expression analysis of differentially expressed genes in blood samples isolated from NM1 WT and KO mice. (a) Gene ontology analysis based on all differentially expressed genes between experimental conditions shows biological process-, cellular component-, and KEGG pathway-associated gene ontology terms plotted in descending order based on their significance. (b) Gene ontology analysis based on only upregulated genes in NM1 KO samples. (c) Heatmap of all differentially expressed genes between NM1 WT and KO blood samples associated with the GO term “Blood coagulation”. (d) Heatmap of all differentially expressed genes between NM1 WT and KO blood samples associated with the GO term “Platelet activation”.

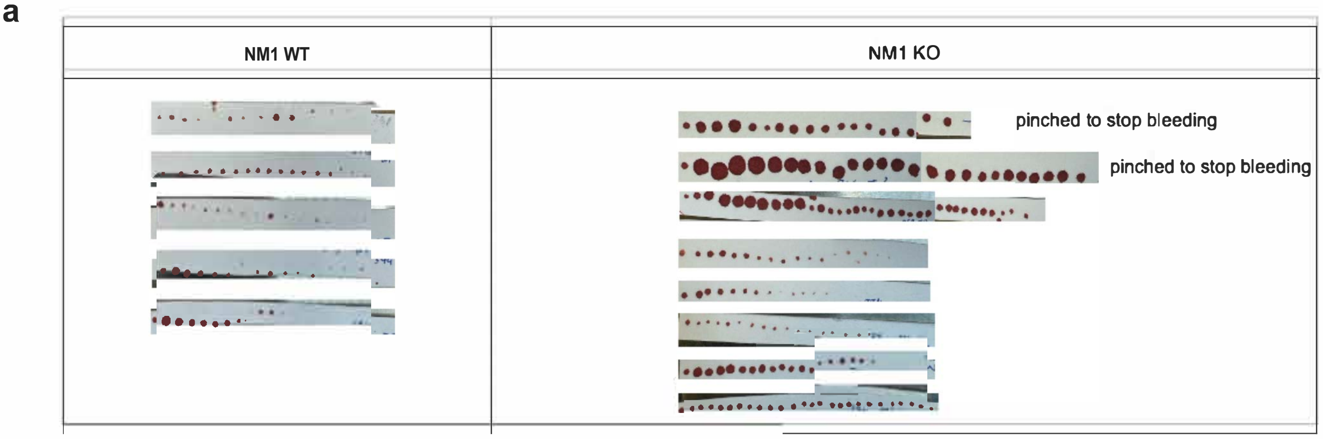


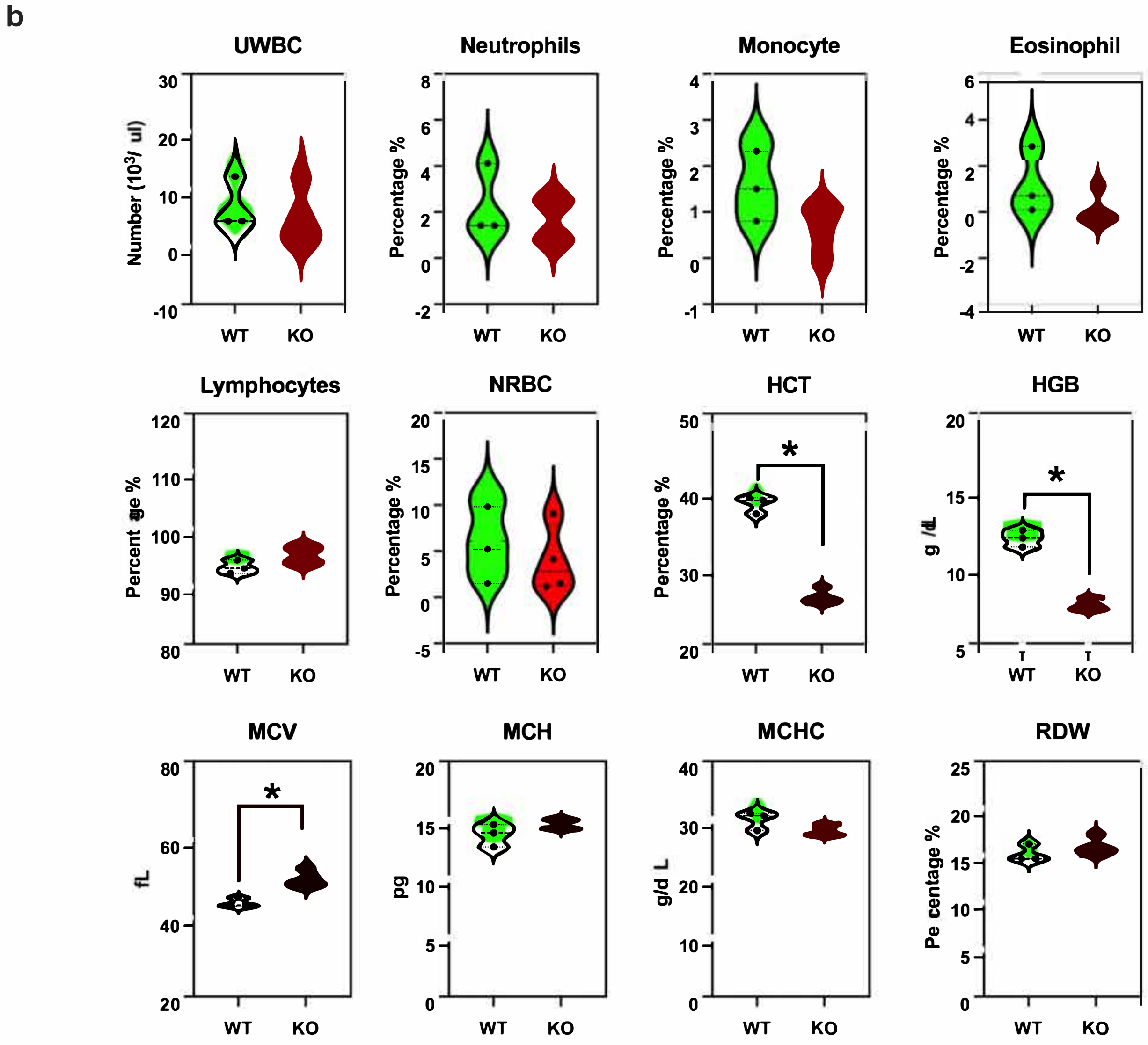


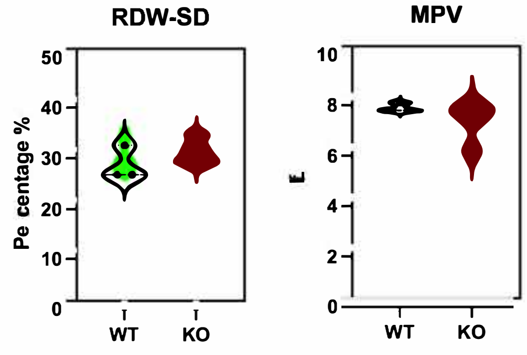


Supplementary figure 3. CBC profiles in WT and NM1 KO mice. (A) Image of blotting paper with blood drops collected from the bleeding time experiment, showcasing the procedure used for measuring bleeding time in both WT and KO mice. (B) Violin plots illustrating additional CBC profiles, including UWBC, differential WBC counts (neutrophils, lymphocytes, monocytes, eosinophils, basophils), NRBC, HCT, HGB, MCV, MCH, MCHC, RDW, RDW-SD, and MPV for WT and KO mice (n = 3 WT, 4 KO). The two-tailed Mann-Whitney U test was used for statistical analysis. *$p < 0.05$.

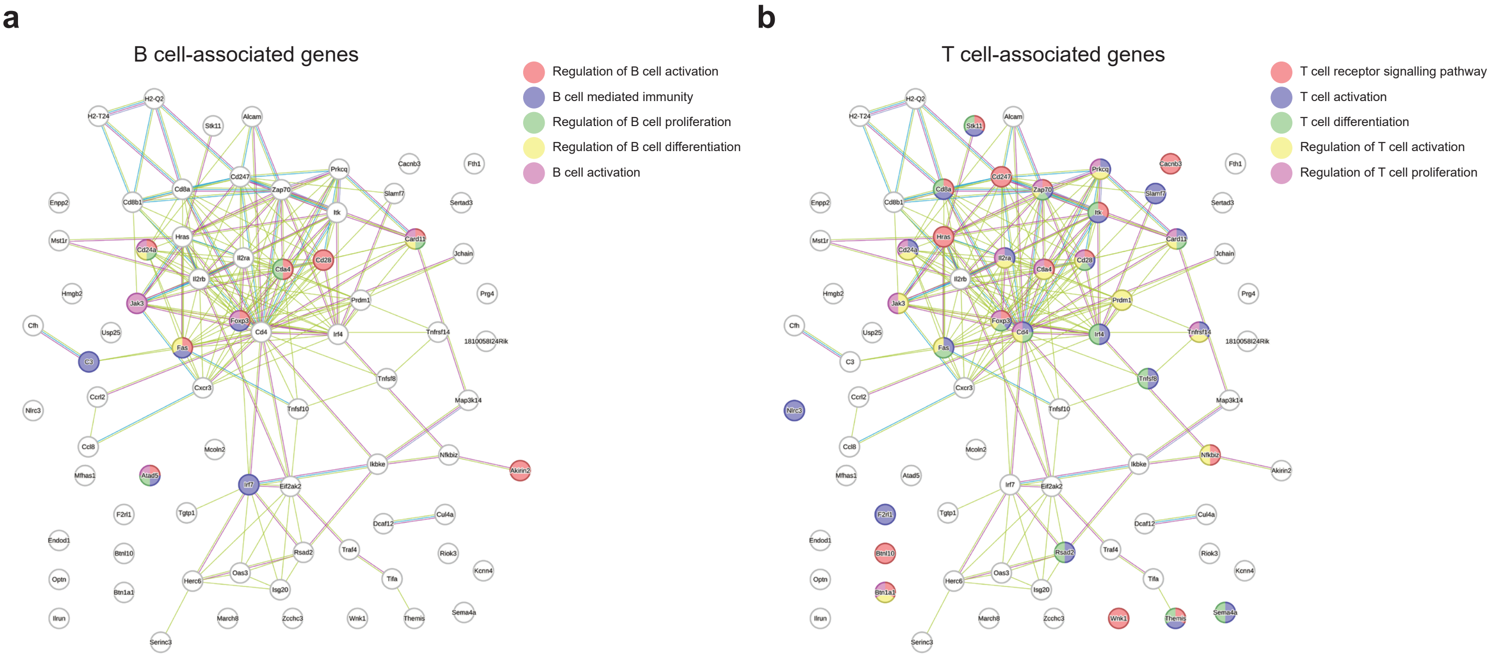


Supplementary Figure 4. STRING analysis of differentially expressed genes associated with B-cell and T-cell biology. (A) String analysis diagram of all genes associated with GO term "Immune system". Colored genes represent the B cell-associated genes differentiated according to the top 5 GO terms related to B cell biology. (B) String analysis diagram of all genes associated with GO term "Immune system". Colored genes represent the T cell-associated genes differentiated according to the top 5 GO terms related to T cell biology.
